# IN4MER Bioink: A Phosphorescent Biosensing Bio-ink for Multiple Analytes (Glucose, Lactate, Oxygen) Measurements and Temperature Sensing Applications

**DOI:** 10.1101/2025.04.22.650078

**Authors:** Waqas Saleem, David Chimene, Berkley P. White, Kaivalya A. Deo, Jeremy L. Thomas, Nithin Chidambara, Cole Mandrona, Kirsten Landsgaard, Brian Ko, Roland Kaunas, Akhilesh K. Gaharwar, Michael J. McShane

## Abstract

3D bioprinting has revolutionized tissue engineering by enabling researchers to create much more complex structures than was practical with earlier techniques. Bioprinting uses computer-controlled layer-by-layer deposition of a mixture of hydrogels and living cells and the resulting structures can mimic the complex geometries of many living tissues by incorporating multiple bioinks with varied material properties and cell populations, allowing researchers to design structures that vary not only in shape, but also in mechanical, chemical, and biological properties throughout the bioprinted construct. However, techniques for evaluating these living constructs and monitoring them over time have not yet caught up to these innovations. Here we describe a novel approach to reporting nutrient values in real-time throughout the scaffold itself, accomplished by dispersing oxygen, glucose, and lactate sensitive microspheres within bioinks. These can be noninvasively interrogated using low-cost phosphorescence lifetime readers to determine and track nutrient concentrations across our bioprinted constructs in real time. The wealth of information this technique produces suggests this may provide a powerful new tool for evaluating and designing future bioprinted constructs.

## 1. Introduction

3D bioprinting involves the successive layer-by-layer deposition of a mixture composed of a biocompatible hydrogel matrix, live cells, and biochemical cues – collectively referred to as *bioink* – which is printed in a specific user-defined pattern to form a scaffold structure.[1] By regulating the physiological and biochemical milieu of the scaffold, the resulting microarchitecture closely resembles the native extracellular matrix (ECM) environment.[2] A viable bioink formulation must also possess favorable rheological properties toward cell viability and printing applications. Most bioinks developed thus far use polymeric hydrogels laden with cells. Polymeric materials from different material sources such as polysaccharides [alginate, κ-carrageenan (κCA)], proteins [gelatin or its methacryloyl form (GelMA), albumins], decellularized materials, and synthetic polymers [polyethylene glycol diacrylate (PEGDA)] have been effectively demonstrated towards bioink preparation.[3] Natural polymers provide similarities to the native ECM and possess biocompatible and biodegradable properties along with their intrinsic gelation mechanisms.[4] Also, synthetic nanosilicate clays, such as Laponite, are widely mixed with bioinks owing to their favorable rheological properties for printing applications, such as shear thinning and yield strength behavior.[5] The incorporation of nanocomposites in hydrogels has led to the development of smart hydrogels with tunable physical properties and degradation profiles, allowing homogeneous distribution of cells in a highly hydrated and mechanically supportive environment.[6,7]

The choice of polymeric material and overall rheological properties of any bioink formulation are critical in determining the final fate of the incorporated cells. Beyond certain bioink formulations, which are specifically designed to promote cell adhesion, proliferation, or differentiation and protect cell viability throughout the bioprinting process, a range of methods can further modulate these cellular behavior, including ligand-based modifications (e.g., RGD peptides, laminin-derived peptides, growth factors like EGF and FGF)[8–11], physical and mechanical cues (e.g., scaffold architecture, stiffness, shear stress)[12–14], and co-culture strategies.[15,16] Therefore, researchers often must optimize the bioink formulations specific to the cell type so that the bioink’s viscoelastic properties favor ECM production and organization, which is critical for cell survival. To address the cell needs- and tissue-specific bioink formulations, it is imperative to have appropriately balanced bioink components, such as microstructure, correct ECM choice, and mechanical properties. Given the diversity of cellular/tissue requirements, metabolic needs, and functional complexity and mechanical properties required for the final intended application a one-formulation-fits-all solution is rarely available.

Beyond the bioink composition, it is notable that the diffusivity of metabolites through the scaffold also strongly impacts the overall cell viability, functionality, and cellular processes. Within a 3D-bioprinted scaffold, nutrients and small metabolites (such as O2 and glucose) must diffuse from the surrounding media solution into the hydrogel matrix to nourish the cells within. Once again, the choice of polymeric-matrix material becomes critical here, as it presents a diffusion barrier to these metabolites. The local concentration of the diffused metabolites through bioprinted structure depends on several factors including but not limited to porosity and pore size; and viscosity, composition, and mechanical/rheological properties of the printed bioink.[17,18] For example, after incorporating perfused vascular channels within the scaffold, Eltaher et al. demonstrated a 3D bioprinted construct of a vascularized human nose that otherwise would have been necrotic.[19] Similarly, increased porosity, pore size, and connectivity of pores can influence cell adhesion, migration, and proliferation activities.[20]

Poor distribution of metabolites or structural limitations of the scaffold may create metabolite-deficient zones within the 3D-printed constructs. Understanding the cellular responses under high metabolic demand (such as elevated glucose requirement and hypoxia) further requires monitoring the spatiotemporal distribution of analytes to confirm necessary supply is achieved, otherwise metabolite-deficient zones may develop within the bioprinted constructs. For example, glucose and lactate concentrations are an important metabolite to monitor. Many cells in culture rely on glucose as an energy source and cell growth can be stalled below critical threshold concentrations. Lactate, a byproduct of cellular metabolism, is another key parameter to monitor, as excessive accumulation can be detrimental to cell viability. Furthermore, the ratio of glucose consumption to lactate production can provide insight into the metabolic state of the cells, indicating whether they are undergoing glycolysis or oxidative phosphorylation. As the level of complexity within these constructs increases, it becomes more challenging to perform mapping and distribution analysis of dispersed metabolites. Invasive methods for analyte monitoring, such as needle-like insertion probes, pierce through the scaffold and provide only localized measurements.[21,22] Alternatively, planar-sensing foils, such as one for O2 monitoring, provide mapping distribution of analyte limited to the contact interfaces.[23–25]

Previous advancements in multianalyte biosensing have primarily focused on concurrent detection of glucose and lactate alongside uric acid or oxygen, or oxygen with ethanol.[26–29] Initial efforts employed electrochemical methods with micro-electrodes/-needles or screen printed electrode platforms.[26,27,30] These later transitioned to optical detection using photoluminescence and fluorescence intensity sensing. While these studies paved the way for standalone multianalyte biosensors towards comprehensive analyte measurements, examples of real-time monitoring under live cell culture conditions remain scarce.[27]

Recently, researchers addressed this gap by incorporating various optical-sensing moieties with diverse sensing modalities into bioink formulations. This approach enabled the investigation of spatiotemporal distribution of different analytes within bioprinted constructs. However, these studies were limited to single metabolite monitoring such as oxygen, adenosine, and dopamine molecules.[31–33]

This work aims to bridge the existing gaps in optical-based multianalyte biosensing within 3D bioprinted structures. We propose a novel approach that leverages the incorporation of discrete microscale biosensing moieties directly into the biomaterial ink formulations. This strategy facilitates seamless metabolite monitoring through a single-mode (phosphorescence) measurement system, overcoming limitations of previous methods. The microparticles are designed with polyelectrolyte nanofilm coatings deposited via a layer-by-layer self-assembly method; we have reported previously that these self-assembled nanofilm coatings provide tunability for the diffusion control of the target analyte and co-reactants into a given microparticle. Moreover, the polymeric matrices in which these microparticles were dispersed had the least impact on the biosensing output.[34–38] For bioprinting, we explore dispersing our biosensing microparticles into various biomaterial ink formulations belonging to diverse polymeric sources to endow them with biosensing features.

In this study, we dispersed optically responsive, oxygen-, glucose-, lactate- and temperature-sensitive microparticles within various 3D bioprinting ink formulations (**Figure 1**). Optical responses in terms of phosphorescence lifetimes were recorded from the hydrogel sensors under exposure to varying analyte concentrations to assess biosensing performance. Ink formulations from three diverse sources, alginate or κCA, GelMA, and PEGDA, were chosen to demonstrate the versatility of biosensing microparticles. Biosensing ink formulations were also 3D bioprinted into constructs and tested against non-printed sensors. To evaluate potential for vivo deployment, 3D bioprinted GelMA glucose sensors were implanted under the subcutaneous tissue of a pig animal, and sensor responses were collected for four months, followed by histology analysis. Finally, biosensors fabricated from bio-ink formulation were used under cell culture conditions for real-time monitoring of O2, glucose, lactate and temperature. The findings demonstrate the effective combination of biosensing particles in biomaterials for oxygen, glucose and lactate biosensing along with temperature monitoring using different ink formulations, the biocompatible nature of 3D-printed constructs, sensitive multianalyte responses under *in vitro* cell culture conditions, and the *in vivo* application of 3D-printed constructs towards use with implanted devices.

**Figure 1:**
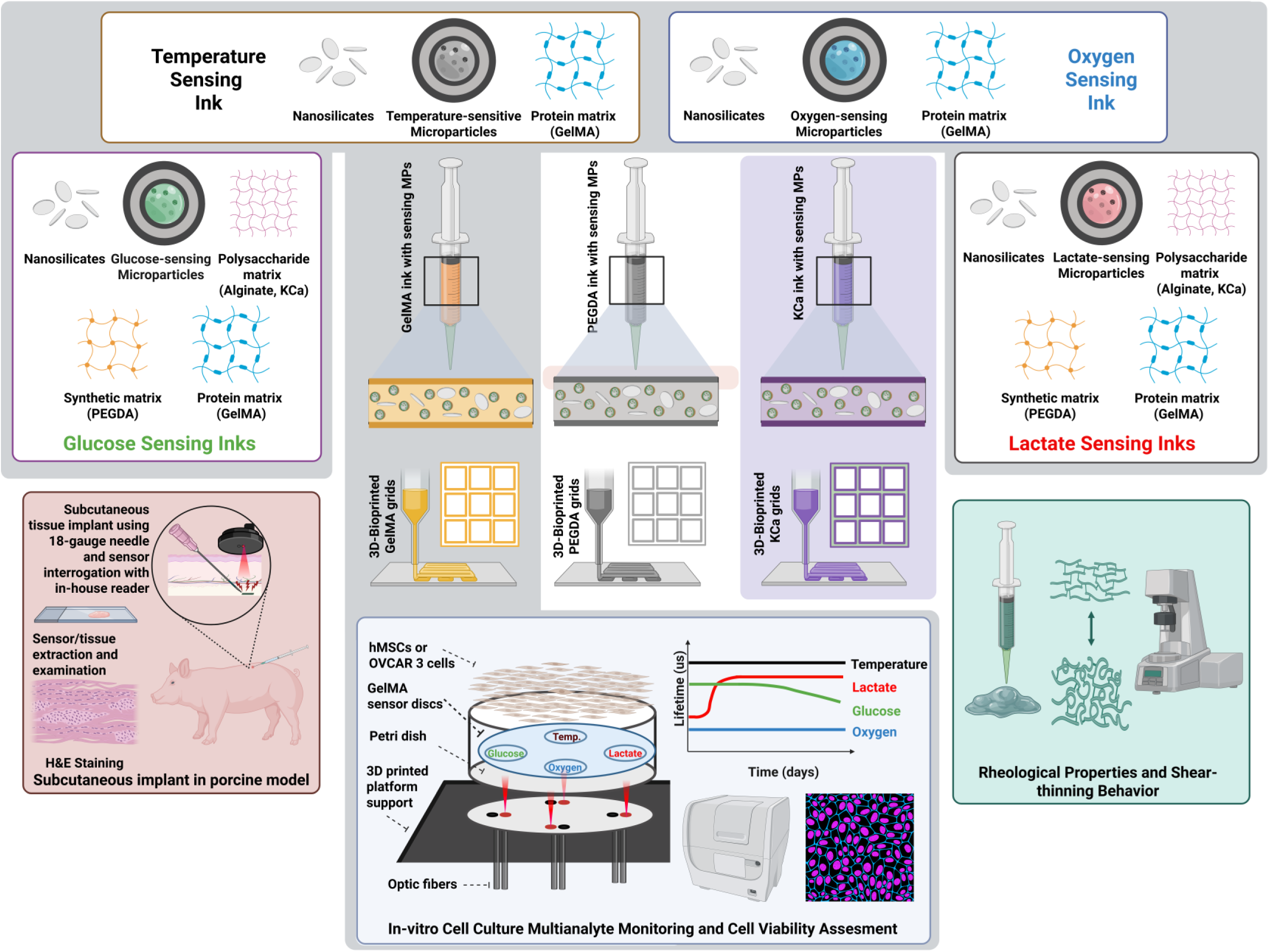
An overview of IN4MER biosensing bioink platform components with demonstrated multidisciplinary applications

## 2. Results and Discussion

This study aimed to transform traditional bioinks into biosensing-enabled formulations. We evaluated the optical responses and rheological properties of various biomaterial inks derived from diverse polymers, incorporating glucose- and lactate-sensing microparticles, respectively. **Supplemental Figure S1** outlines the overall approach for developing glucose-sensing microparticles, employing our previously reported strategies and findings from other similar work. [36,38–40] The fabrication process involved embedding the oxygen-sensitive dye Palladium(II) meso-tetra(sulfophenyl)tetrabenzoporphyrin (HULK) and oxidase enzymes (glucose oxidase and catalase) within alginate microparticles. Subsequently, varying numbers of self-assembled layers of oppositely charged polyelectrolyte nanofilms were deposited onto the microparticle surface using the layer-by-layer (LbL) self-assembly method. As demonstrated in our previous work[36,40], these polyelectrolyte nanofilm coatings are critical for controlling the diffusion of the target analyte molecules such as glucose and lactate within the alginate microparticle core while oxygen transport remains significantly unaffected[38], thereby providing an additional means to optimize sensor sensitivity and dynamic range. To optimize the number of LbLs and microparticle concentration, we fabricated various iterations of monolithic slab gels within an alginate matrix. Sensor responses were then evaluated to establish the “reference MPs” and “reference gel” for glucose sensing. Additionally, lactate-sensing microparticles were formulated; it is noteworthy that they had lower enzyme content (six times lower activity) compared to the glucose-sensing particles. For these, we strategically employed a water-soluble composite form of PdBP dye encapsulated in ethylcellulose nanoparticles (ECNP) due to its superior sensitivity and dynamic range compared to free HULK dye, as reported in our previous work[38,41], a step taken to enhance the sensitivity and dynamic range for the lower enzyme levels. We leveraged our knowledge of lactate-sensing microparticles, employing 30 LbL coatings and a 1X microparticle concentration as the Reference MPs. Similarly, the temperature-sensing PAN particles (oxygen insensitive) were fabricated as described in our previously published work.[42] Briefly, these microparticles are inherently insensitive to oxygen changes, but are sensitive towards temperature changes. All of the phosphor forms incorporated into the microparticles (HULK, ECNP, and BMAP) are red excitable (630 nm) in nature, with long emission wavelengths in near-infrared region (800-810 nm); see **Supplemental Figure S2** for excitation and emission scan spectra from individual sensing materials.

To demonstrate the versatility of our biosensing approach, the reference glucose- and lactate-sensing microparticles were incorporated into diverse bioink formulations derived from various sources such as carbohydrates [к-carrageenan (κCA)], proteins [methacrylated-gelatin (GelMA)], synthetic polymers [polyethylene glycol dimethacrylate (PEG-DA)] and nanosilicates (Laponite XLG). We first evaluated each blank ink formulation (without MPs) using a custom-built 3D bioprinter to assess printability. We employed a 400 μm nozzle (22-gauge taper tip) at a flow rate of 0.15 mL/min (printing speed of 20 mm/s). These parameters were chosen based on previously published results using murine 3T3 preosteoblast cells to ensure optimal resolution, fidelity, and minimal stress on embedded cells.[6]

Our goal was to establish bioprintable inks suitable for incorporating microparticles. We systematically evaluated various combinations of minimal polymer gels and shear-thinning biomaterial, i.e., Laponite XLG (**Supplemental Figure S3**), to identify formulations with decent printability and resolution. Based on qualitative analysis, promising blank inks were down-selected: 6% GelMA with 1% Laponite XLG, 20% PEGDA with 4% Laponite XLG, and 1% κCA with 1% Laponite XLG. These blank inks were then incorporated with either glucose-sensing (**Figure 2 and 3**) or lactate-sensing (**Figure 4 and 5**) reference MPs, respectively. We fabricated both monolithic gel slabs and 3D bioprinted structures using these formulations. For evaluation, circular punches (from slabs) or cut strips (from bioprinted grids) were obtained from each crosslinked formulation. Samples were assessed for oxygen and either glucose or lactate sensing responses using the testing setup described in **Supplemental Figure S4**. We compared these responses within each sample (intra-sample) and against reference gels (inter-sample) to investigate the impact of 3D printing on sensor performance (**Table 1**). Combining composition optimization, microparticle inclusion, and 3D bioprinting, this comprehensive approach establishes a foundation for developing advanced bioprinted sensors with customized functionalities.

**Figure 2:**
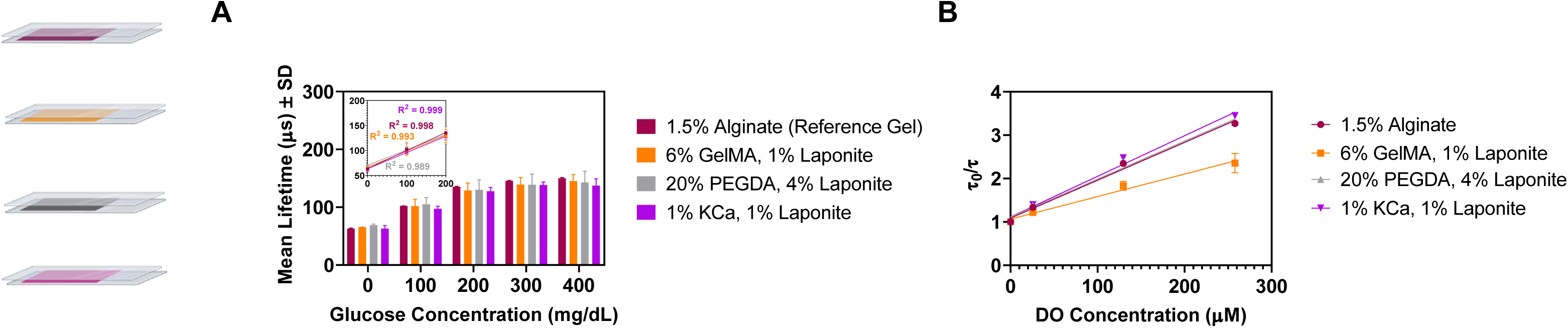
**(A)** Overlaid glucose response plots of glucose sensors (N=3) from slabs showing linear response to glucose challenges up to 400 mg/dL of analyte exposure (with linear portion of responses shown within inset graph). Two-way ANOVA indicated no significant differences between mean lifetimes across different samples when exposed upto 400 mg/dL of glucose concentration **(B)** Overlaid Stern-Volmer plots from glucose sensors (N=4) fabricated from gel slabs made with reference MPs dispersed in different matrices.

**Figure 3:**
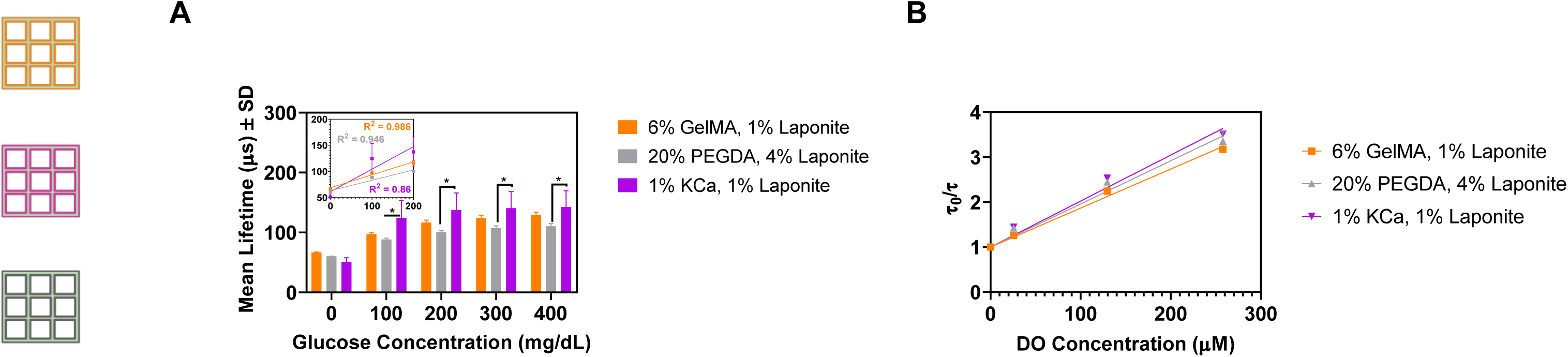
**(A)** Overlaid glucose response plots of glucose sensors (N=3) from 3D-printed gird sensors showing linear response to glucose challenges up to 400 mg/dL of analyte exposure (with linear portion of responses shown within inset graph). Two-way ANOVA indicated significant differences between mean lifetimes values of PEGDA vs KCa gels only for 100, 200, 300 and 400 mg/dL of glucose concentrations. (* = p < 0.05) **(B)** Overlaid Stern-Volmer plots from glucose sensors (N=4) fabricated from 3D-printed grid samples with reference MPs dispersed in different matrices.

**Figure 4:**
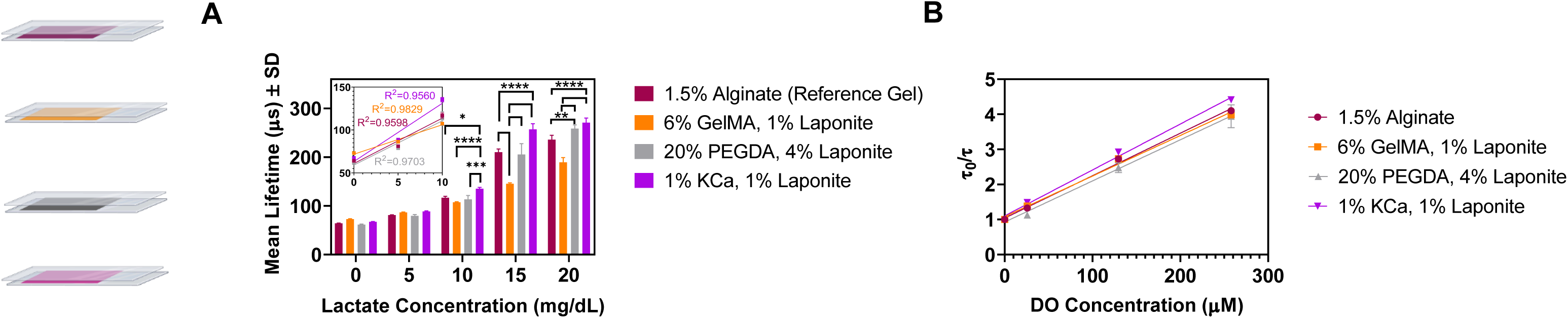
**(A)** Overlaid lactate responses plot of lactate sensors (N=3) from slabs showing linear response to lactate challenges up to 20 mg/dL of exposure. Two-way ANOVA indicated no significant differences between mean lifetimes of any given gel upto 5 mg/dL lactate concentration. However, significant differences were observed between Alginate vs. KCa, GelMA vs. KCa, and PEGDA vs. KCa gels at 10 mg/dL; Alginate vs. GelMA/KCa, GelMA vs. PEGDA/KCa at 15 mg/dL; Alginate vs. PEGDA/KCa, and GelMA vs. PEGDA/KCa at 20 mg/dL of lactate concentration, respectively. (* = p < 0.05, ** = p < 0.005, *** = p < 0.0005, **** = p < 0.0001) **(B)** Overlaid Stern-Volmer plots from lactate sensors (N=4) fabricated from gel slabs in different matrices.

**Figure 5:**
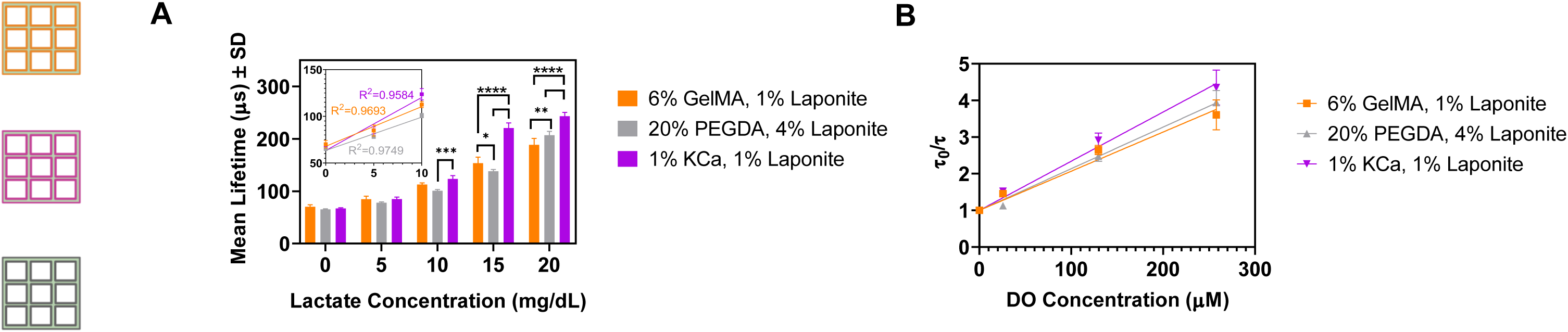
**(A)** Overlaid lactate responses plot of lactate sensors (N=3) from 3D-printed grid sensors showing linear response to lactate challenges up to 20 mg/dL of exposure. Two-way ANOVA indicated no significant differences between mean mean lifetimes values for any given 3DP gels formulation upto 5 mg/dL. Significant difference were observed between PEGDA vs. KCa gels at 10 mg/dL; GelMA vs. PEGDA/KCa and PEGDA vs. KCa at 15 mg/dL of lactate concentration; GelMA vs. PEGDA/KCa and PeGDA vs. KCa gels at 20 mg/dL of lactate concentrations, respectively. (* = p < 0.05, ** = p < 0.005, *** = p < 0.0005, **** = p < 0.0001) **(C)** Overlaid Stern-Volmer plots from lactate sensors (N=4) fabricated from 3D-printed grid samples in different matrices.

**Table 1.** Summary of obtained calculated values from various form-factor of glucose and lactate sensors in varying ink formulations.

| Samples | $K_{sv}$<br>[$\mu\text{M}^{-1}$]<sup>a)</sup> | LOD [mgdL<sup>-1</sup>]<sup>b)</sup> | MDGC or MDLC<br>[mgdL<sup>-1</sup>]<sup>c)</sup> | Dynamic range<br>[mgdL<sup>-1</sup>] | Sensitivity<br>[us.dL.mg<sup>-1</sup>]<sup>d)</sup> |
| --- | --- | --- | --- | --- | --- |
| Circular Discs [3 x 0.75 mm]<sup>e)</sup> |  |  |  |  |  |
| AnA-Ref gel | 0.0093 | 2.1 | 139.9 | 137.8 | 0.58 |
| κCA | 0.0099 | 41.5 | 116.6 | 75.1 | 0.99 |
| GelMA | 0.0083 | 8.3 | 227.7 | 219.4 | 0.40 |
| PEGDA | 0.0099 | 13.1 | 185.5 | 172.4 | 0.48 |
| 3DP-Strips [4 x 0.4 x 0.4 mm]<sup>f)</sup> |  |  |  |  |  |
| κCA | 0.0102 | 4.3 | 82.1 | 77.8 | 1.18 |
| GelMA | 0.0087 | 0.8 | 179.8 | 179.0 | 0.35 |
| PEGDA | 0.0096 | 2.5 | 170.1 | 167.5 | 0.30 |
| Circular Discs [3 x 0.75 mm]<sup>g)</sup> |  |  |  |  |  |
| AnA-Ref gel | 0.012 | 1.1 | 9.3 | 8.2 | 6.44 |
| κCA | 0.014 | 0.8 | 9.5 | 8.7 | 7.80 |
| GelMA | 0.012 | 1.6 | 6.7 | 5.2 | 6.44 |
| PEGDA | 0.011 | 1.1 | 5.2 | 4.1 | 12.64 |
| 3DP-Strips [4 x 0.4 x 0.4 mm]<sup>h)</sup> |  |  |  |  |  |
| κCA | 0.013 | 1.5 | 7.4 | 5.9 | 9.64 |
| GelMA | 0.011 | 1.4 | 7.1 | 5.7 | 5.60 |
| PEGDA | 0.011 | 1.8 | 9.0 | 7.3 | 4.93 |

<sup>a)</sup> Stern-Volmer constant; <sup>b)</sup> Limit of detection; <sup>c)</sup> Maximum detectable analyte (glucose or lactate) concentration; <sup>d)</sup> From 0 to 400 mgdL<sup>-1</sup> (glucose) or 10 mgdL<sup>-1</sup> (lactate) sensor, respectively; <sup>e, f)</sup> Glucose sensors; <sup>g, h)</sup> Lactate sensors.

Rheological measurements, including peak hold, strain sweep, and yield stress determination, were performed on selected biomaterial ink formulations both with and without reference MPs (**Table 2**). These tests were aimed at assessing the printability and material properties of the inks.

**Table 2.**
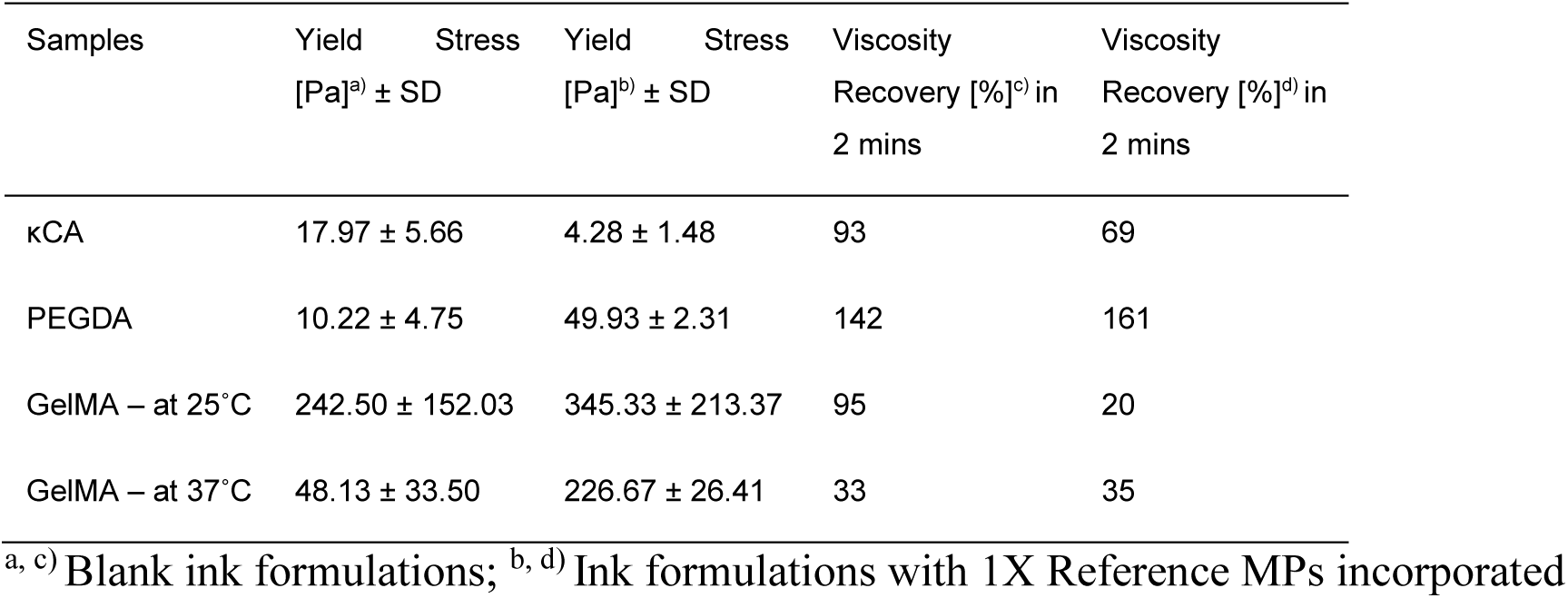
Summary of rheological assessments obtained from blank and MPs-containing ink formulations.

Following rheological characterization, *in vivo* biocompatibility assessments were conducted using 3D-bioprinted GelMA strips subcutaneously implanted in a porcine model (**Figure 6**). This *in vivo* evaluation provided valuable insights into the safety and potential for long-term applications of the developed bioinks. Finally, the GelMA-based ink formulation was used to fabricate four different types of biosensors, each sensitive to oxygen, glucose, lactate and temperature, respectively, followed by real-time simultaneous multi-sensor monitoring under *in vitro* cell culture conditions (**Figure 7 and 8**) and cytocompatibility assessments.

**Figure 6:**
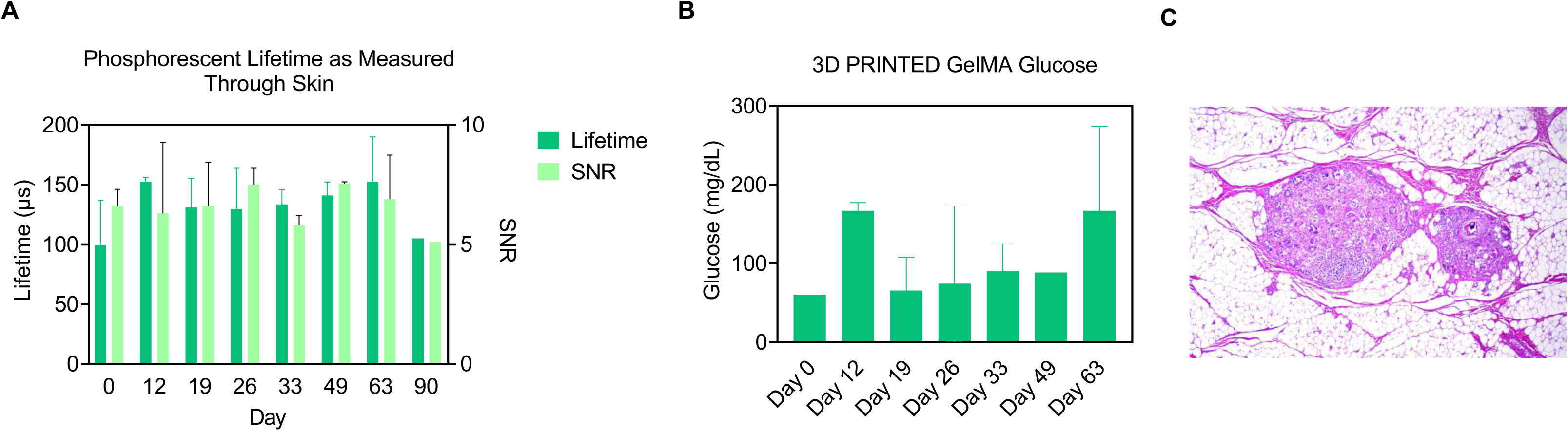
(A) Mean lifetime measurements and SNR values (N=2) from two separate 3D bioprinted GelMA glucose biosensor implants over the course of 3 months. (B) Back-calculated glucose concentration values from 3D bioprinted GelMA glucose biosensors obtained through *in vitro* calibration results. (C) Zoomed section of tissue with sensor demonstrating expected biodegradability, minimal capsule formation, and tissue healing through chronic phagocytosis.

**Figure 7:**
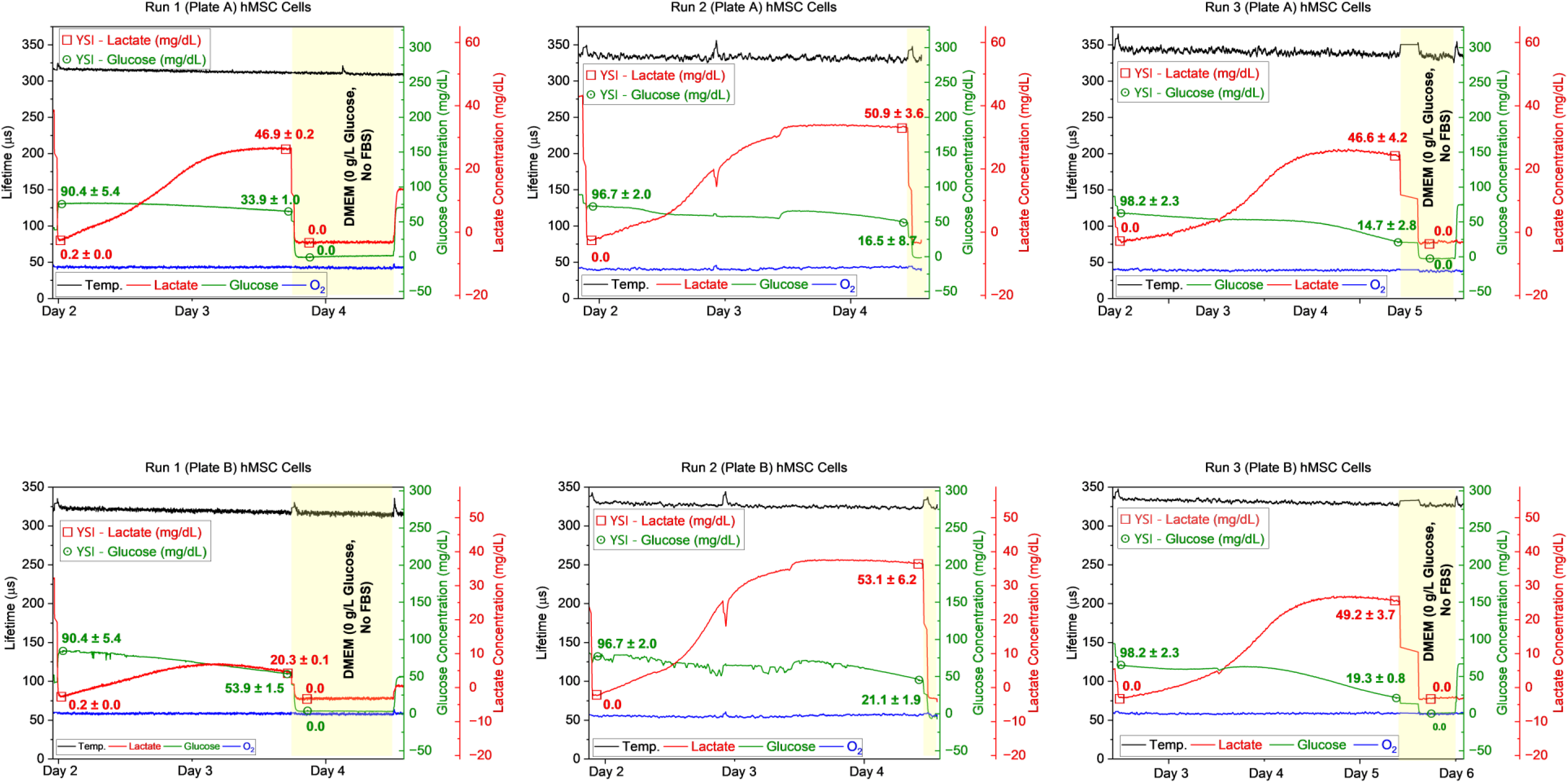
Multianalyte sensors from GelMA-based ink formulations seeded with 0.2 million cells/ml of induced pluripotent cell derived mesenchymal stem cells (ihMSCs) showed lactate production (∼5-35 mg/dL from sensors) and glucose depletion changes.

**Figure 8:**
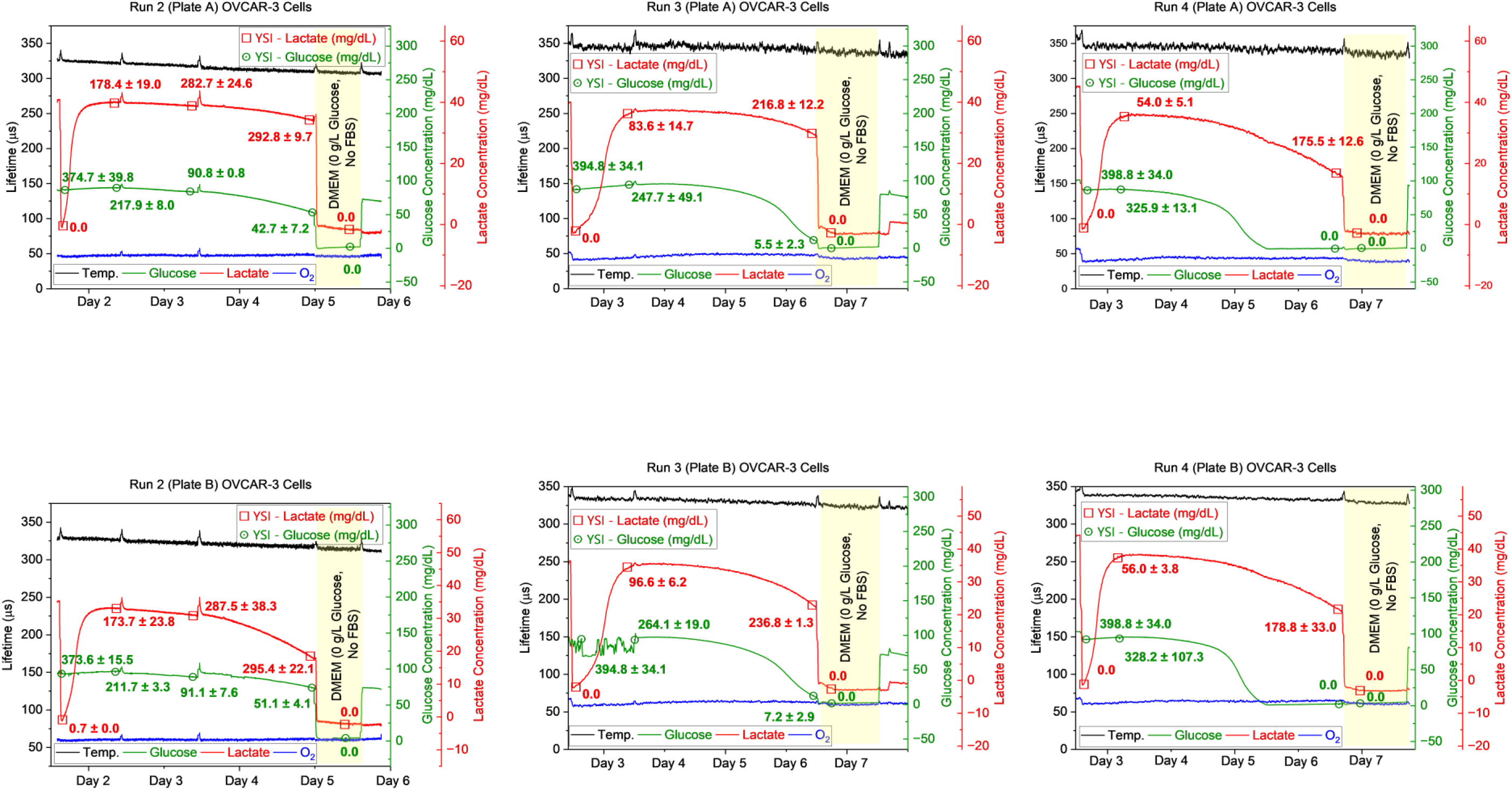
Multianalyte sensors from GelMA-based ink formulations seeded with 0.4 million cells/ml ovarian cancer cells (OVCAR-3) showed robust spike in lactate production (>20 mg/dL) and glucose depletion changes.

### 2.1. Interplay between LbLs, MPs Concentration, and Sensor Response

We investigated various formulations for glucose-sensing alginate microparticles with varying numbers of LbL coatings (5-40) and microparticle concentrations [11 mg/ml (0.5X), 22 mg/ml (1X), 44 mg/ml (2X)] within a calcium-crosslinked alginate matrix. Each formulation was thoroughly evaluated (N ≥ 3) using monolithic hydrogel slabs. These slabs underwent sequential testing: first, their responses to increasing glucose concentrations (up to 400 mg/dL) and batch-to-batch variability of glucose-sensing microparticles assessment [**Supplemental Figure S5(A-C)]** followed by oxygen sensitivity responses [**Figure S6(A-B)**].

Based on the comprehensive data, microparticles with 35 LbL coatings at 1X concentration exhibited the most desirable characteristics for glucose sensing. These microparticles demonstrated a highly linear response with stepwise changes in response to increasing glucose levels, achieving an R-squared (R^2^) value of 0.9477 for the entire range [**Supplemental Figure S5(A)**]. Similarly, for lactate sensing, microparticles with 30 LbL coatings and a 1X concentration displayed the most linear response to increasing lactate concentrations (up to 20 mg/dL).

To further analyze the impact of LbL coatings and microparticle concentration on sensor performance, a 3D plot was generated, visualizing the relationship between these variables and the corresponding R^2^ value from glucose calibration plots [**Supplementary Figure S5(B)**]. This analysis revealed an interdependent relationship between these factors.

Generally, increasing the number of LbL coatings creates a diffusion barrier at the microparticle surface, hindering the diffusion of analytes (except oxygen) to the core. This can serve as a user-adjustable parameter for optimizing sensor’s sensitivity, linearity and dynamic range. The studies also emphasize the importance of MP concentration, as the overall amount of incorporated microparticles can influence sensor response. While higher microparticle concentration often leads to improved linearity, it can also compromise the sensor’s dynamic range.

### 2.2. Oxygen Sensitivity Responses and Batch-to-batch Reproducibility

The oxygen sensitivity responses from the glucose-sensing and lactate-sensing gels were separately assessed. Stern-Volmer (SV) plots were generated to calculate SV-constant (*Ksv*) values to assess oxygen sensitivity. We performed comprehensive oxygen sensitivity response testing over various combinations of alginate gels with varying LbLs and MP concentrations for glucose-sensing microparticles embedded with free HULK dye. As represented in **Supplemental Figure S6(A-B)**, the 35 LbLs at 1X MPs concentration from glucose-sensing microparticles resulted in the most linearized quenching behavior with decreasing oxygen concentrations from 21% (∼257 µM DO) to 0% (0 µM DO) air, resulting in one of the highest *Ksv* values (0.009 µM^-1^). The higher sensitivity and *Ksv* value (0.012 µM^-1^) were reported for lactate-sensing gels compared to glucose-sensing gels; this behavior was attributed to using the PdBP dye impregnated into ethylcellulose polymeric nanoparticles in the lactate sensing particles, as noted above and as shown in **Supplemental Figure S7(A)**.

The batch-to-batch reproducibility of glucose-sensing reference microparticles (35 LbLs, 1X concentration) was assessed by testing sensors from two separate batches with increasing glucose concentrations (0-400 mg/dL). **Supplemental Figure S5(C)** shows the linear region of lifetime changes for glucose concentrations up to 200 mg/dL. Statistical analysis confirmed no significant difference in the mean lifetime responses between sensors from both batches, demonstrating consistent sensing performances.

Following this, both glucose and lactate reference microparticles were individually dispersed in various biomaterial ink formulations. Their oxygen sensitivity and analyte responses were then compared to their corresponding reference gels.

#### 2.2.1. Oxygen sensitivity and glucose-sensing responses from Glucose Reference MPs in diverse polymeric matrices

Glucose-sensing Reference MPs were individually dispersed in 6% GelMA with 1% Laponite XLG, 20% PEGDA with 4% Laponite XLG and 1% κCA with 1% Laponite XLG ink formulations, respectively, and monolithic gel slabs were fabricated. Circular discs (N≥3) were punched from the slab and tested under incremental glucose challenges, and responses were compared against the Reference Gel (**Figure 2A**). Later, oxygen sensitivity responses from the same discs were also measured, and SV plots (**Figure 2B**) were generated to determine *Ksv* values. Based on our extensive experience and premise that the diffusion control of the target analyte inside a given microparticle is significantly governed by the thickness of nanofilms coated over the microparticle surface and not by the dispersed matrix, we expected no significant differences in terms of glucose-challenge responses and oxygen sensitivity differences between various ink formulations dispersed with the glucose Reference MPs.[38] As expected, there were no statistically significant differences between responses from various gels when exposed to incremental glucose challenges up to 200 mg/dL. All hydrogels provided a comparable oxygen sensitivity response, with 1% κCA with 1% Laponite XLG providing the highest *Ksv* value (0.0099 µM^-1^).

Separately, the same ink formulations were utilized for 3D bioprinting grid structures, subsequently crosslinked, and later tested against glucose challenge and oxygen sensitivity responses [**Figure 3(A-B)**]. Here, except for κCA samples, the 3D-printed samples from GelMA and PEGDA ink formulations provided no significant differences in glucose sensing responses. In contrast, there were significant differences between κCA vs. GElMA and PEGDA at 0 mg/dL, respectively, and κCA against PEGDA ink formulations. Comparatively, 3D bioprinted strips (grids) provided a lower sensitivity response against circular discs (slabs), which can be explained in terms of the differences between the size and dimension of tested samples [circular discs: 3 mm (dia) X 0.75 mm (h), strips: 3 mm (l) X 800 m (dia)]. **Supplemental Figure S3(B)** depicts stereotaxic images of 1” x 1” bioprinted grid (3 x 3) structures (two layers) premixed with glucose Reference MPs in all three diverse polymeric matrices.

Table 1 summarizes the oxygen sensitivity of the hydrogel sensor punches, expressed as Stern-Volmer constants (*KSV*), which were calculated using Equation[43] (1) based on sample lifetimes recorded at 10 s intervals under varying oxygen levels[44]:

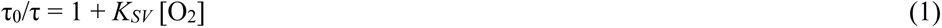

where *KSV =* Stern-Volmer constant, **τ**_0_ = lifetime under no dissolved oxygen concentration, **τ** = lifetime under given dissolved oxygen concentration, and [O2] = dissolved oxygen concentration, respectively. Other critical parameters such as limit of detection (LOD), Maximum Detectable Glucose Concentration (MDGC), dynamic range and sensitivity from non-printed and 3D printed sensors fabricated from all ink formulations are also summarized in **Table 1**. For both non-printed and 3D bioprinted (3DP) sensors, except for κCA gels, oxygen sensitivity of all gels in terms of *KSV* values were comparable, with similar LODs, and a dynamic range of at least 137 mg/dL. Differences in the sensitivities among non-printed and printed ink formulations can be attributed to the different dimensions and sizes (surface area, volume) of the tested sensors and bioprinting process for fabrication. Similarly, except for κCA gels, the detection limit for non-printed and printed sensors was the lowest at 2.1 mg/dL for alginate gels (non-printed) and 0.8 mg/dL for GelMA gels (3DP). Finally, the maximum detectable glucose concentration (MDGC) was provided by GelMA gels (non-printed and 3D-bioprinted).

One explanation for the observed variations in κCA gel properties could be due to differences in crosslinking and resulting changes in mechanical strength. κCA gels rely on ionic crosslinking via hydrogen bonds, facilitated by high molarity KCl solutions (5% in our experiments). These potassium ions interact with the negative charges within the κCA polymer chains, allowing adjacent hydroxyl groups to form intermolecular hydrogen bonds. Studies have shown that efflux or replacement of potassium ions with other divalent cations can weaken these bonds, reducing gel strength.[45] In our experiments, replacing 10 mM CaCl2 with 0.1 M KCl in TRIS buffers for analyte challenges might not have provided sufficient KCl concentration to maintain optimal gel strength under flow conditions. In conclusion, GelMA gels exhibited more consistent response behavior and were less affected by sensor dimensional variations.

#### 2.2.2. Oxygen sensitivity and lactate-sensing responses from Lactate Reference MPs in diverse polymeric matrices

To establish a truly multifunctional and multianalyte biosensing platform technology for traditional bioinks, we expanded one step further to incorporate lactate-sensing reference MPs into down-selected printable ink formulations from diverse polymeric sources.

A similar approach was followed, wherein lactate-sensing Reference MPs were first dispersed into alginate matrices, crosslinked into the monolithic slab, and tested for lactate sensing and oxygen sensitivity responses, thereby establishing baseline response of the lactate Reference Gels [**Supplemental Figure S7(B)**]. Later, these lactate reference MPs were individually dispersed into various printable blank ink formulations. Monolithic slabs and 3D bioprinted grid structures were then separately fabricated and crosslinked before testing. The individual material responses to the lactate challenge and oxygen sensitivity were subsequently determined (**Figure 4 and 5**).

Except for κCA formulation, lactate sensors from monolithic slabs revealed no significant difference in mean lifetime responses between alginate against GelMA and PEGDA ink formulations [**Figure 4(A)**] when tested at 0 mg/dL lactate. At 10 mg/dL lactate challenges, there were also significant differences among mean lifetime responses between alginate vs. GelMA and PEGDA formulations. In contrast, no differences were observed between GelMA vs. PEGDA, and κCA vs. alginate, GelMA and PEGDA formulations. Similarly, at 20 mg/dL lactate challenges, significant differences were observed between alginate vs. GelMA and PEGDA, and κCA vs GelMA and PEGDA formulations. Again, no significant differences were observed between GelMA and PEGDA formulations. **Figure 4(B)** depicts the overlaid Stern-Volmer plot from all four gel formulations, where oxygen sensitivity responses in terms of *Ksv* values (**Table 1**) from all gels (except for κCA gels; being slightly higher) were comparable.

A trend from 3D bioprinted lactate-sensing materials was observed, similar to glucose-sensing materials, where no significant differences in mean lifetime responses were observed at 0 mg/dL of lactate challenges across all ink formulations [**Figure 5(A)**]. With subsequent incremental challenges of 10 and 20 mg/dL of lactate, respectively, significant differences in mean lifetime responses were only observed by κCA against GelMA and PEGDA formulations. Similarly, **Figure 5(B)** depicts overlaid Stern-Volmer plots from all three formulations of 3D bioprinted sensors revealed similar oxygen sensitivity responses (same *Ksv* values) for GelMA and PEGDA sensors, but slightly higher for κCA gels.

**Supplemental Figure S3(C)** presents stereotaxic images of lactate sensing 3D bioprinted grid structures from all three gel formulations. As summarized in **Table 1**, also similar to glucose sensors, all gel formulations for lactate sensors exhibited comparable oxygen sensitivity—except κCA gels. Non-printed alginate gels and 3D-printed GelMA gels demonstrated the lowest LODs at 2.1 mg/dL and 0.8 mg/dL, respectively. All sensors exhibited a dynamic range exceeding 4 mg/dL. While sensitivity varied among formulations, GelMA-based sensors displayed consistent performance across both non-printed and 3D-bioprinted configurations, highlighting their potential for *in vitro* lactate monitoring applications.

One possible explanation for the higher variability in responses between different formulations for lactate-sensing can be attributed to the limited amount and low total enzyme activity of the enzyme loaded during the microparticle fabrication process. Because the specific activity of native lactate oxidase is relatively low, and incorporating enzyme mass beyond a certain threshold presents fabrication challenges, lactate reference microparticles (MPs) are typically fabricated with a five-fold lower total enzyme activity (∼ 1000 U) compared to glucose reference MPs (> 6000 U). We hypothesize that these response variabilities arise from insufficient enzyme concentration and total enzyme activity and can be mitigated through optimized enzyme loading in future studies.

### 2.3. Rheological behavior of biosensing ink formulations

We evaluated the impact of microparticle addition on the rheological properties of the biomaterial inks by dispersing glucose-sensing Reference MPs in various formulations and comparing them to their blank counterparts. Shear-rate sweep tests [**Supplemental Figure S8(A)**] revealed that all tested formulations retained their shear-thinning behavior, even after microparticle incorporation. κCA inks displayed minimal changes in shear-thinning properties, while PEGDA inks containing microparticles exhibited a higher viscosity compared to blank formulations. GelMA inks, being thermally responsive, underwent rheological testing at two crucial temperatures (25°C and 37°C) corresponding to points above and below the gelation point. At 37°C where the viscosity is lower, MP-containing GelMA inks exhibited increased viscosity – roughly one order of magnitude – compared to their blank counterparts. In contrast, minimal viscosity changes were observed after addition of microparticles when testing at 25°C due to the overall increased viscosity (∼1-2 orders of magnitude) relative to the blank gels.

Building upon shear-rate sweep results, peak hold tests were conducted to simulate viscosity changes under varied shear rates. Non-thermoresponsive ink formulations (κCA and PEGDA) were evaluated at 25°C, while thermoresponsive GelMA ink formulations were assessed at both 25°C and 37°C **[Supplemental Figure S8(B)]**. At 25°C, blank PEGDA exhibited the most rapid viscosity recovery, exceeding its baseline value by 142%, followed by GelMA (95%) and κCA (93%). However, when microparticles were incorporated, only PEGDA maintained rapid viscosity recovery (161%), while κCA (69%) and GelMA (20%) exhibited significantly reduced recovery.

For GelMA inks at 37°C, both blank and microparticle-containing formulations displayed lower viscosity recovery, reaching only 33% and 35%, respectively. While the microparticle-containing GelMA ink showed slightly higher viscosity, both formulations demonstrated slower and less complete viscosity recovery compared to the blank GelMA at this temperature.

To better understand and simulate bioprinting conditions with thermoresponsive materials like GelMA, a separate peak hold test was conducted. This test mimicked the temperature shifts experienced by the bioink as it transitions from the warm syringe barrel (37°C) to the cooler room temperature (25°C), as depicted in **Supplemental Figure S8(C).** Moreover, it replicated the shear rate variations encountered within the syringe (wall shear), extruder tip, and finally, the printed structure (low shear). This allowed for the evaluation of viscosity changes during the bioprinting process, accounting for both shear rate variations and temperature fluctuations as the bioink transitions from 37°C within the syringe barrel to 25°C at or outside the extruder tip.

As expected, the blank GelMA bioink demonstrated significant thermal responsiveness and shear-thinning properties, leading to an effective reduction in shear stress during the extrusion process.[6] It exhibited rapid viscosity recovery, exceeding its baseline value prior to the gel-crosslinking step. At 37°C, GelMA inks exhibit lower viscosity, mimicking a liquid-like state that minimizes extrusion shear stress and facilitates smooth printing. In contrast, the GelMA bioink with microparticles, while also exhibiting a similar recovery behavior, demonstrated significantly slower and lower recovery compared to the blank formulation.

To further assess rheological properties, we performed strain sweeps to determine changes in storage (*G’*) and loss (*G’’*) moduli before and after the inclusion of microparticles [**Supplemental Figure S8(D)**]. For κCA gels, the blank ink exhibited stable elastic behavior with a higher *G’*, which subsequently decreased after the addition of microparticles, leading to a lower *G’* possibly due to the disruption of polymeric side-chain interactions by microparticles required for crosslinking of κCA gels. *G’’* increased with increasing strain for both formulations, suggesting a viscous-like behavior. Similarly, the *G’* of GelMA inks containing MPs was higher at both temperatures compared to their blank counterparts, suggesting gel strengthening after microparticle inclusion. However, at 37°C, the GelMA with MPs ink formulation appeared to enhance strain-thinning behavior and promote transition towards a more viscous behavior compared to blank inks at both temperatures. Finally, with PEGDA inks, the addition of microparticles demonstrated a limited effect on the storage and loss modulus properties of the ink.

To conclude the rheological assessment, the dynamic stress sweep method was employed to determine the yield stress of all ink formulations, with results presented in **Supplemental Figure S8(E)**. Consistent with the previous tests, GelMA exhibited the highest yield stress at both tested temperatures, further increasing with microparticle inclusion. Statistical analysis revealed no significant differences in yield stress between blank and MP-containing GelMA at either temperature. In contrast, significant differences were observed for PEGDA and κCA inks: microparticle incorporation strengthened the PEGDA gel, while it reduced the gel strength of κCA inks. These findings further emphasize the material-specific impact of microparticle addition on yield stress, highlighting the importance of considering material properties when selecting and incorporating microparticles for bioprinting applications.

### 2.4. *In Vivo* response of 3D bioprinted glucose sensors

Because bioprinted constructs are often implanted to support regeneration of tissue, we set out to gather data on the long-term *in vivo* effects of microsphere-containing scaffolds on surrounding tissue. For this experiment, we chose glucose-sensitive microparticles embedded in the GelMA ink formulation. GelMA was chosen because it is enzymatically biodegradable. GelMA ink formulations containing 1X glucose-sensitive microparticles were 3D bioprinted into 5 mm long scaffold strips using a 22-gauge taper tip needle (2 layers). Subsequently, the bioprinted scaffolds were subjected to electron beam (E-beam) irradiation at a dose < 25 kGy. Based on our previous experiments with similar scaffold material[41], we predicted that a GelMA-based scaffold would be partially degraded over the 3 months of this experiment, allowing for interactions between microspheres and the immune system. No cells were included in the inserted scaffolds.

Five (5 mm ✕ 800 µm) scaffolds were inserted just under the skin of a healthy, immunocompetent Yorkshire Cross female. Insertion sites were intentionally varied to cover different skin thicknesses, from the thickest skin at the shoulder to the much thinner skin of the ribs and flanks. These samples were interrogated using a 1D reader head to excite the phosphors through the skin using red light, while the lifetime of the escaping near-infrared phosphorescence was quantified using an incorporated photodetector [**Supplemental Figure S9(A)**]. Scaffolds were tested regularly after insertion, starting a few hours after the procedure. In line with our previous *in vivo* tests, scaffolds could not be reliably located and read immediately after insertion; this depends on several factors but is likely primarily a consequence of local inflammation caused by the insertion procedure.[41] However, by the second measurement point (day 12), consistent readings could be collected; this persisted for the rest of the experiment. This suggests that the inflammation response to implantation subsided after the healing process and did not drive a chronic negative reaction.

As expected, the samples producing the most reliable readings were the two located under the thinner rib skin. These samples were determined to be a little over 5mm from the surface at their shallowest points. Readings for these scaffolds (A8 and H9) [**Supplemental Figure S9(A-B)**] showed consistent readings averaging 132 ± 20 μs and 142 ± 23.60 μs, respectively (**Figure 6A**). Signal to noise ratios averaged 5.95 ± 1.19 and 7.24 ± 1.36, respectively. These lifetimes fall comfortably within the expected ranges established by the earlier *in vitro* experiments, indicating that the microspheres continued reporting local glucose concentrations as expected throughout the experiment. In parallel, in vitro calibration studies were conducted **[Supplemental Figure S9(D)]** using a separate cohort of post E-beam irradiated sensors from the same batch as those implanted in vivo. These studies demonstrated a lack of responsiveness to glucose **(Figure 6B)**, likely attributed to the use of glucose-sensing microparticles with non-crosslinked nanofilms. This post-fabrication step of nanofilm crosslinking with glutaraldehyde is crucial for enabling sensor operation under low-oxygen (30-40 µM) conditions, as it significantly reduces glucose diffusion (179%) while minimally affecting oxygen transport. Despite this limitation, the successful 3D printing and *in vivo* detection of sensors with comparable lifetimes suggest promise for further development of implantable biosensors, especially considering the implanted sensors were not fully optimized for functioning under low-oxygen tissue microenvironment conditions.

Biocompatibility studies are crucial for evaluating new materials that interact with living cells. We aimed firstly to evaluate the scaffold’s effects on the body in terms of host interface and healing response. Secondly, we evaluated the body’s effects on the scaffold, quantifying the device’s state after 3 months *in vivo* and testing how long we would be able to detect a signal from the embedded scaffold after insertion. Therefore, independent veterinary pathologists were tasked with evaluating the scaffold’s insertion site for scaffold presence, state, skin zone, host interface, and healing response.

At gross examination, the examined scaffold was found to still be visible within the subcutis – just over 5mm from the epidermal surface – and could still be readily detected using the reader head [**Supplemental Figure S9(E)**]. After sectioning and staining, histologic evaluation revealed the scaffold undergoing remodeling classified as partial infiltrative, as macrophages and multinucleated giant cells could be seen containing device fragments throughout the scaffold [**Figure 6C and Supplemental Figure S9(F)**]. This is evidence of the ongoing chronic phagocytic biodegradation of the inserted scaffold. Capsule formation was minimal. There was no evidence of further response to the scaffolds in the surrounding local tissue, supporting the existing *in vitro* evidence of bio- and cytocompatibility.

Overall, these *in vivo* experiments served as a pilot study to evaluate the performance and biocompatibility of bioprinted scaffolds containing microspheres in the body. We established that the scaffolds were detectable through skin comparable to human skin and could be detected *in vivo* throughout the 3-month experiment. Histologic evaluation showed that the scaffolds were undergoing remodeling via chronic phagocytosis with barely discernible capsule formation, and no evidence of further response or toxicity in the surrounding tissue. These promising results suggest that the biosensing microspheres may be adapted to applications involving implantable scaffolds. However, given the small scale of these experiments, a more extensive study with more samples and subjects will be necessary to draw broader conclusions.

### 2.5. Multianalyte real-time biosensing under cell culture conditions

For proof-of-concept demonstrations of multianalyte biosensing under in vitro cell culture conditions, monolithic sensor slabs were fabricated using a GelMA-based ink formulation. Circular sensor discs were then punched out from these slabs and subsequently sterilized by E-beam irradiation at a dose < 25 kGy. The choice of GelMA was intended to promote cell growth and infiltration while assessing whether the sensors affected normal cell growth. The circular sensor design enabled effective optical response data collection, ensuring adequate coverage of both excitation and emission sources from the optical fiber, while maintaining consistent dimensions across experiments. We selected a relatively high seeding density of 200,000 cells/ml in a 60 mm petri dish in order to measure detectable metabolite changes in a 2-3 day experimental time span. MSCs are known to exhibit a relatively low rate of glucose consumption in vitro— approximately 98 fmol/cell/h, corresponding to about 0.0112 mg per 10^6 cells per hour. Based on this rate, an estimated 0.2688 mg of glucose is consumed over a 24-hour period. To ensure that the change in glucose levels was sufficiently large for accurate measurement, a relatively high-density culture was employed.[46] This setup balances the risk of experimental limitations related to high analyte concentrations or low cell populations.

#### 2.5.1. Real-time cell culture monitoring of multianalytes with non-transformed cell line

A general procedure described in **Supplemental Figure S10 (A-B)** was used for *in vitro* cell culture experiments and multi-sensor setup for data collection purposes. As anticipated, the multianalyte biosensors used with ihMSCs showed minimal to no changes in lifetime responses for oxygen and temperature sensors throughout different sets of the experiments (**Figure 7**). Glucose sensor lifetime responses exhibited a gradual decline, characterized by an initial period of low glucose consumption followed by a progressively increasing depletion rate after 2–3 days. Notably, glucose concentration measurements from Run 1 (Plate B) and Run 3 (Plates A and B) strongly correlated with concurrent YSI glucose measurements. Conversely, the remaining glucose sensors deviated from YSI measurements by ±20 mg/dL. Lactate sensors, in contrast, demonstrated a gradual increase in lifetime responses, indicative of sustained lactate production. With the exception of the lactate sensor in Run 1 (Plate B), all other lactate sensors showed substantial lactate production, particularly near or beyond Day 4. At these time points, lifetime changes from most lactate sensors approached peak lifetime values of 200–235 µs, corresponding to the maximum detection threshold of 20 mg/dL of lactate concentration [**(Supplemental Figure S12 (A)**]. YSI lactate measurements at these peak points yielded concentrations of ∼40–50 mg/dL, slightly higher but reasonably close to the sensor readings of 30–35 mg/dL. It is important to note that the limited dynamic range (0–20 mg/dL) of our lactate sensors precluded accurate measurement of lactate concentrations exceeding 20 mg/dL (or lifetime values >200 µs). This limitation stems from the combination of the low specific activity of the commercially-sourced lactate oxidase as well as the low enzyme loading during the emulsification step. These factors constrained our optimization efforts for lactate sensors within the scope of this proof-of-concept study. Future work will focus on enhancing the sensitivity and dynamic range of these lactate-sensing micro-particles for broader applications.

#### 2.5.2. Real-time cell culture monitoring of multianalytes with transformed cell line

A similar trend in multianalyte biosensing responses was observed with ovarian cancer cells (OVCAR-3), where minimal to no changes were noted for temperature and oxygen sensors (**Figure 8**). However, lactate sensors revealed a significant increase in lactate production, as indicated by a rapid surge in lifetime responses. We expected high lactate production from OVCAR-3 cells, given that cancer cells are known to overproduce lactate especially under hypoxia. The rate and extent of lactate production in OVCAR-3 cells compared to ihMSCs was of particular interest.[47–49] E-beam irradiated glucose sensors in DMEM high glucose complete media yielded lifetime measurements of approximately 150 µs (corresponding to a glucose concentration of ∼100 mg/dL), significantly lower than YSI glucose measurements, which ranged from ∼375– 400 mg/dL. This difference is attributable to the inherently lower sensitivity (100 µs total bandwidth) of our non-e-beam irradiated glucose sensors across the 0–400 mg/dL dynamic range. Post-e-beam irradiation, the glucose sensor sensitivity remained at 100 µs total, but within a narrowed dynamic range of 0–100 mg/dL **[Supplemental Figure S12(A)]**. As glucose depletion progressed during cell culture (post-Day 2), approaching 100 mg/dL, agreement between calculated glucose sensor concentrations and YSI glucose measurements improved. The observed increase in lactate sensor lifetimes (approaching ∼250 µs) indicated complete lactate sensor saturation at approximately 30–35 mg/dL, while YSI lactate measurements were substantially higher (2.5–10-fold). Following these peak lifetime ranges, all lactate sensors exhibited a consistent decline in lifetime values, indirectly suggesting a reduction in lactate production. However, this decline in lifetime output was completely discordant with YSI lactate measurements, which, conversely, showed a trend of consistently increasing lactate production. The most probable explanation for this discrepancy is a significant decrease in catalytic activity[50] or conformational changes to the enzyme’s active site[51], potentially due to the lower acidic environment. Lactate oxidase reportedly exhibits limited stability and a pH dependence (pH 6.0) for optimal catalytic activity. Furthermore, a mutant lactate oxidase from *Aerococcus viridans* was specifically bioengineered by Garcia-Morales et al. for optimal function under low acidic conditions. Further experiments are needed to elucidate these effects, which were beyond the scope of this study. Nonetheless, our proof-of-concept demonstrates a unique method for visualizing the Warburg effect[52–54] in real-time, highlighting the potential of our approach for investigating metabolic profile changes under stressed conditions.

##### Cytocompatibility Assessments

Live/dead staining coupled with confocal fluorescence microscopy revealed that the majority of cells remained viable, particularly at the center of the dish where both ihMSCs and OVCAR-3 cells were equidistant from the sensors. In contrast, a zone of low cell density was consistently observed at the periphery of the glucose sensors (Figure S11). This reduced cytocompatibility is hypothesized to result from the continuous activation of the glucose sensors—driven by the persistent high concentrations of glucose (100–450 mg/dL) in the culture medium—which may lead to localized depletion of glucose and oxygen. Such conditions could create a stressed microenvironment that limits cellular attachment in close proximity to the sensor in the first place. Notably, viable cells were observed adhering to the surfaces of both lactate and temperature sensors, and in vivo histological analysis of the bioprinted glucose sensors demonstrated biocompatibility and cellular infiltration. These observations suggest that the adverse effects observed at the periphery of the glucose sensors may be attributable to their continuous activity, in contrast to lactate sensors that become active only upon lactate production. Further investigations are warranted to elucidate the underlying mechanisms, although such studies are beyond the scope of the present work.

## 3. Conclusion

In this study, we explored the integration of biosensing functionalities into a range of bioink formulations derived from different polymeric sources using a simple ratio-mixing approach. We successfully incorporated various analyte-sensing and temperature-sensing microparticles, demonstrating the interplay between microparticle concentration and nanofilm thickness on sensor sensitivity, dynamic response range, and linearity. Our findings confirmed that the biosensing properties of the bioinks were primarily determined by the incorporated microparticles rather than the native polymeric matrices.

The experiments validated that 3D bioprinted and non-printed sensors from different bioink formulations provided consistent sensor responses, particularly at lower analyte concentrations. Notably, we observed significant differences between the κCA formulation and the GelMA and PEGDA formulations, attributable to differences in crosslinking mechanisms. Rheological assessments indicated minimal effects of microparticle inclusion on shear-thinning behavior and recoverability, while mechanical strength variations provided insights into sensor response variability.

Our *in vivo* studies further established the utility of 3D bioprinting for the fabrication of implantable hydrogel-based glucose sensors. *In vitro* cell culture studies demonstrated the responsiveness of biosensors to changing metabolite levels under higher metabolic demand, reinforcing their potential for future applications. Nevertheless, the operational flexibility of these optically-sensing microparticles and their functionality post-Ebeam irradiation support future applications in bioreactor and metabolic profile monitoring setups.

In future studies, we plan to incorporate and optimize the use of a Phosphorescence Lifetime Imager (PLI) system.[42] The PLI system provides 2D images of real-time phosphorescence intensity changes from analyte-sensitive microparticles dispersed within the hydrogel scaffold, enabling direct visualization of analyte distribution and consumption in real-time. Ultimately, the optimized PLI system would facilitate multiplexed analyte monitoring within the 3D bioprinted scaffold, thus enabling the realization of our ultimate goal: spatio-temporal mapping and tracking of analyte distribution within the scaffold. Additionally, we aim to expand our array of biosensors to detect other relevant metabolites, such as amino acids and ketone bodies, thereby enriching our understanding of local metabolites in cell culture and biomedical research.

## 4. Experimental Section/Methods

### Materials

The following chemicals were purchased from specified suppliers for use in the experiments: Laponite XLG was purchased from BYK Additives, sodium alginate (CAS No. A215875000 - MW: 100000 Da), dimethyl sulfoxide (CAS No. 67-68-5), calcium chloride (CAS No. 22350), calcium carbonate (CAS. No. 239216), TRIS base (CAS No. T1503), and catalase (3000-5000 U/mg) from bovine liver (CAS. No.C9322) from Sigma-Aldrich, Inc., St. Louis, MO; polyethylene glycol diacrylate (PEG-DA, 3400 MW)from Thermo Fisher Scientific, Waltham, MA; collagen (rat tail-type I) and neutralization solutions from Advanced Biomatrix, Carlsbad, CA; Trizma hydrochloride (TRIS HCl) (CAS No. VWRB85827), and phosphate buffer saline (PBS) from VWR Chemicals LLC, NY, USA; glucose oxidase (76 U/mg) from Aspergillus niger (CAS No. G0050), (Z, Z, Z)-Sorbitan tri-9-octadecenoate (SPAN 85) (CAS. No. S0064), Polyoxyethylenesorbitan Trioleate (TWEEN 85) (CAS. No. T0547) from Tokyo Chemical Company Co., LTD, Tokyo, Japan; lactate oxidase II (Prod # L1175, 1KU) from AG Scientific phosphor dye as palladium (II) meso-tetra-(sulfophenyl) tetrabenzoporphyrin sodium salt (HULK dye; CAS No. T41161 - MW: 1327.55 g/mol), Palladium (II) meso-tetra-(4-carboxyphenyl) tetrabenzoporphyrin (PdBP dye; Cat No. T13343 - MW 1095.41 g/mol) from Frontier Specialty Chemicals, Logan, UT; isooctane (CAS No. 94701) from Avantor Performance Materials, LLC, Randor, PA. All chemicals were used without further purification. Tetrahydrofuran (THF) (CAS No. 109-99-9) and poly(sodium-4-styrenesulfonate) (PSS) (CAS No. 25704-18-1, average MW 70 kDa) were obtained from Sigma Aldrich. Acetone (CAS No. 67-64-1) was obtained from Macron Fine Chemicals. Poly(vinylidene chloride-co-acrylonitrile) (PViCl-PAN) (CAS No. 9010-76-8) was obtained from Polysciences, Inc. Palladium (II) tetramethacrylated benzoporphyrin (PtBMAPP) was donated by PROFUSA, Inc. For cell culture applications, 10% (v/v) fetal bovine serum (FBS) was purchased from Atlanta Biologicals. Gibco’s Dialyzed FBS (Cat # A3382001) and Dulbecco’s Modified Eagle Medium (DMEM) high glucose, were purchased from Thermo Fisher Scientific. α-minimum essential media from Invitrogen, 4 mM L-glutamine, and 100 U/mL penicillin and streptomycin were procured from Sigma Aldrich.

### Glucose-sensing microparticle fabrication

Monodisperse alginate microparticles of sizes (10±1 microns) were synthesized using the water-in-oil emulsion method. Briefly, a 500 µL volume of 10 mM (in DMSO) phosphor dye was added to 5 mL of 3% aqueous solution of sodium alginate. Nutate the solution overnight at room temperature. The next day, 81.9 mg of glucose oxidase and 91.5 mg of catalase enzymes were premixed in 2 mL of Tris-HCl buffer (pH 7.2; w/o. CaCl2), transferred to the alginate-HULK dye mixture, and nutated for 20 mins. For emulsification, a 50 mL centrifuge tube filled with 10.8 mL of isooctane solution (premixed with 290 µL of Span 85 surfactant) was spun at 8000 rpm speed using an IKA T25 homogenizer. To this solution, the alginate-HULK-enzymes cocktail solution was slowly added and mixed for 10 seconds, followed by the addition of 1.5 mL of isooctane solution mixed with 130 µL of Tween 85 surfactant. The solution was mixed for 10 seconds, followed by a last addition of 4 mL of 10% CaCl2 solution and mixed for another 15 seconds. The mixture to transferred to a round bottom flask with a magnetic stirrer and stirred at 300 rpm (20 mins) for ionotropic gelation. The mixture was centrifuged in a 50 mL centrifuge tube and spun at 627 RCF for 2 mins; the supernatant was discarded and washed 3X in DI water at 157 RCF for 1 min. The supernatant was discarded, and microparticles and layer-by-layer self-assembly of polyelectrolyte nanofilms coating procedure was carried out.

### Lactate-sensing microparticle fabrication

Lactate-sensing alginate microparticles were fabricated using our previously published method with slight modifications.[38] First, we replaced free HULK dye with a water-soluble composite form of Palladium (II) benzoporphyrin (PdBP) dye encapsulated in ethylcellulose polymeric nanoparticles. The polymeric modification of dye has several advantages over free dye, such as higher sensitivity changes under lower oxygen conditions and aggregation-free dye encapsulation due to an overall lower concentration of PdBP incorporation. A detailed procedure for synthesizing ethyl cellulose nanoparticle suspension solution and microparticle fabrication is described elsewhere.[38] Second, we reduced the total volume reaction for emulsification by five times, but added the entire mass of lactate enzyme (∼18.4 mg; 1000U total), thereby, increasing the enzyme loading efficiency within microparticles. Briefly, 0.75 mL of 4% alginate and 0.25 mL of nanoparticle suspension (1 mg) were mixed for 30 min. Separately, lactate oxidase (entire mass from the vial; 1000 U) and catalase (18.3 mg; >64,000 U) were dissolved in 0.5 mL of 50 mM TRIS buffer and added to the mixture, followed by 20 min nutation. This precursor was then emulsified similarly using 2.16 mL of iso-octane (premixed with 68.4 mg of SPAN 85), 0.3 mL of iso-octane with 29.2 mg of TWEEN 85, and 0.8 mL of 10% w/w CaCl2 solutions. The final emulsion was stirred for 20 min, centrifuged, the supernatant discarded, and microparticles proceeded to layer-by-layer self-assembly of polyelectrolyte nanofilms coating procedure.

### Oxygen-sensing microparticle fabrication

Alginate microparticles capable of sensing oxygen were fabricated using a water-in-oil emulsion method. First, a solution of sodium alginate (3% aqueous) was mixed with HULK phosphor dye (10 mM) and stirred overnight. This mixture was then emulsified with 10.8 mL of isooctane (premixed with 260uL of SPAN 85 surfactant) using homogenizer by spinning at 8000 rpm for 10 seconds, followed by another addition of 1.5 mL of iso-octane (premixed with 130 µL of TWEEN 85 surfactant). Solution was mixed for 10 seconds, and then final addition of 4 mL of 10% CaCl2 solution was carried out and mixed for another 15 seconds. Final emulsified solution was transferred to a round bottom flask and stirred at 300 rpm (20 mins) for ionotropic gelation. The mixture was centrifuged at 627 RCF for 2 mins; the supernatant was discarded and washed 3X in DI water at 157 RCF for 1 min. The supernatant was discarded, and microparticles and layer-by-layer self-assembly of polyelectrolyte nanofilms (5 bilayers) coating procedure was carried out.

### Temperature-sensing microparticle fabrication

To encapsulate the porphyrin dye (PtBMAPP) within polyacrylonitrile (PAN) beads, a mixture of 8.9 mL tetrahydrofuran (THF) and 71.1 mL acetone was first stirred in a glass beaker. Then, 400 mg of poly(vinyl chloride)-polyacrylonitrile (PViCl-PAN), 0.2 mg of PtBMAPP dye, and 1 mL of 1 mg/mL poly(sodium styrene sulfonate) (PSS) solution were added and stirred at a moderate rate until all components dissolved completely. Subsequently, 240 mL of deionized water slowly dripped into the solution while stirring. The covered beaker was then left mixed overnight at 85°C for solvent evaporation. Purification involved adding another 1 mL of 1 mg/mL PSS solution and mixing for 1 hour, followed by centrifugation at 10,000 RCF for 8.5 minutes. The supernatant was discarded, and the pellet was resuspended in deionized water. This centrifugation and resuspension process was repeated to remove impurities, with the final resuspension being in 70% ethanol for two wash cycles. The final purified beads were obtained after a final resuspension and storage in deionized water.

### Layer-by-layer Self-Assembly procedure

The layer-by-layer self-assembly method used here is adapted from our previous publications. [38] Briefly, the microparticles underwent a series of consecutive centrifugation steps in 2% w/w polyallylamine hydrochloride (PAH; pH 8.0) followed by 2% w/w polystyrene sulfonate (PSS; pH 7.2) solutions, resulting in the successive buildup and self-assembly of alternating charges of nanofilms coating over the microparticles surfaces.[55] Here, a single PAH layer over a PSS layer is considered one bilayer, whereas we applied a total of 5, 10, 15, 20, 25, 30, 35, and 40 bilayers, respectively. Microparticles at any given bilayer step were stored in 10 mM Tris-HCl buffer (pH 7.2) supplemented with 10 mM CaCl2 at 4°C.

### Microparticles Characterization

A known volume (30 µL, in triplicates) of microparticles at each bilayer step was taken out and dried overnight in a vacuum oven to determine its dried weight. The determination of microparticle weights enabled the quantification of the concentration of dried microparticles within the initial stock suspension of microparticles. Subsequently, a size distribution analysis of alginate microparticles was assessed using the Nexcelom Cellometer Mini device, which reported successive growth in average microparticle diameter size due to layer-by-layer self-assembly procedure. In addition, the microscopy data indicated minimal aggregate presence within the stock suspension.

### Monolithic Hydrogel Slabs Fabrication

For any given formulation, blank gels were prepared using the same recipe but no reference microparticles were added. The following procedures were followed for reference microparticles containing formulations for hydrogel slabs.

#### 1.5% Alginate gels (Reference gels)

For reference microparticles containing alginate gels (a.k.a. *reference gels*), 8.8 mg of reference microparticles were washed thrice with DI water and resuspended back into a total volume of 75 µL of DI water. To this suspension, 25 µL of 33.3% aqueous suspension of calcium carbonate was added, and the solution was vortexed, followed by 200 µL of 3% w/v alginate solution. The solution was vortexed again, and 100 µL of 50 mM MES buffer (pH 6.1) was added. The mixture was quickly pipetted between two microscope slides sandwiched by a 0.75 mm Teflon space and incubated for 20 mins at room temperature. The acid-mediated dissolution of calcium carbonate led to calcium release that ionically crosslinked the alginate gel. Finally, the gel slab was recovered from the slides, transferred to 10 mM TRIS buffer (with 10 mM CaCl2), and stored at 4℃ for 24 hours before testing.

#### 1% *κ*-carrageenan with 1% Laponite XLG

In a 2mL centrifuge tube, 8.8 mg of reference microparticles were washed thrice with DI water, and the supernatant was removed. 200 µL of 2% w/v *κ*-carrageenan (in DI water) was syringed and added to the centrifuge tube, and the mixture was manually homogenized using a spatula. Finally, 200 µL of 2% Laponite XLG solution was added, and the mixture was vortexed and manually mixed using a spatula for 30 seconds to avoid phase separation issues. The final mixture was collected into the syringe and injected between two microscope slides separated by a 0.75 mm Teflon spacer. The entire assembly was submerged in a 5% KCl solution for 20 minutes, leading to crosslinking adjacent polymeric chains of carrageenan through hydrogen bonds. Finally, the gels were recovered from the slides and stored in a 10 mM TRIS buffer solution with 0.1M KCl.

#### 6% Gelatin-Methacryloyl (GelMA) with 1 % Laponite XLG

Lyophilized GelMA (80% methacrylated) was obtained using a previously established procedure.[6] Briefly, 10 g of gelatin (Bloom No. 300, Type A) was solubilized in 100 mL of phosphate buffer saline (PBS) solution and heated at 60℃ for 1 hour; 8 mL of methacrylic anhydride was added in a dropwise manner over several minutes; solution was maintained at 60℃ for an additional 3 hours followed by addition of 400 mL of PBS; and dialyzed for seven days at 50℃ before final lyophilization. For the final 6% GelMA with 1% Laponite gels containing reference microparticles, 200 µL of 0.5% Irgacure 2959 photoinitiator solution was prepared and briefly sonicated for 60 seconds. Then, 24 mg of lyophilized GelMA was added, and the solution was heated to 55℃ under stir. Once GelMA was completely dissolved, the temperature was brought down to 40℃. Separately, 8.8 mg of reference microparticles in a 2 mL centrifuge tube was washed thrice; the supernatant was removed, then mixed with 200 µL of 2% Laponite XLG solution using a spatula. The Laponite mixture was added to the GelMA mixture at 40℃ and mixed homogeneously for a few minutes. Finally, the mixture was collected through a syringe and injected between two microscope slides, sandwiched by a 0.75 mm Teflon spacer. Slides were irradiated under UV-lamp (dosage: 20 mW/cm^2^) for 2 minutes, and crosslinked gels were recovered and stored in 10 mM TRIS buffer with 10 mM CaCl2 at 4℃ until tested.

#### 20% Polyethylene-glycol Diacrylate (PEGDA) with 4% Laponite XLG

100 mg of PEGDA (MW: 3400) with 150 µL of DI water was added. To this solution, 1.4 mg of water-soluble Irgacure 2959 was added, and the solution was covered from light and nutated for 10 minutes. Separately, 6.6 mg of reference microparticles were washed thrice with DI water and mixed with a 150 µL syringed volume of 8% Laponite XLG solution after the supernatant removal. The total mixture was homogenized manually with the help of a spatula to avoid phase separation issues and then collected in a syringe for injection between two microscope slides separated by a 0.75 mm Teflon spacer. The gel was irradiated under a UV-lamp for 10 minutes (5 mins each side), and the crosslinked gel was transferred to 10 mM TRIS buffer solution with 10 mM CaCl2 at 4℃ for storage until testing.

#### 3D bioprinted grids formation

For all 3D bioprinting applications, a custom-built 3D bioprinter was utilized to print a 1” x 1” grid structure using the following printing parameters: 400 µm size tip (22 gauge taper tip nozzle), printing speed = 10 mm/s), travel speed = 25 mm/s, layer height = 0.4 mm, number of layers = 2, filament diameter = 1.75 mm, extrusion multiplier = 1.5, and infill density = 90%.

#### Polysaccharide ink formulation

Blank polysaccharide ink formulations were prepared by mixing three separate volumes (1 mL ea.) of 2% w/v *κ*-carrageenan, each with 1 mL either 1%, 2%, or 4% w/v Laponite XLG. This resulted in final compositions of 1% *κ*-carrageenan with varying Laponite XLG (0.5%, 1% or 2%), respectively. The blank formulations were filled inside the syringe barrel and 3D printed to form gird structures. After analyzing the printing fidelity and resolution, we down-selected the 1% *κ*-carrageenan with 1% Laponite XLG since it provided the best printing results using lower polymeric compositions. A similar procedure to above for making biosensing ink formulation was followed. First, 44 mg of reference microparticles (based of dried microparticles concentration) was washed thrice with DI water, supernatant discarded, and then mixed with 1 mL of 2% w/v *κ*-carrageenan. The mixture was uniformly mixed using the spatula. Then, 1 mL of 2% w/v Laponite XLG was added to the mixture, homogeneously mixed using a spatula for 2 minutes, vortexed for 30 seconds, and filled inside the syringe barrel to 3D print the grid structure. Finally, printed structures were stored in a 10 mM TRIS buffer (7.2) supplemented with 10 mM CalCl2 at 4℃ until further tests were performed.

#### Protein ink formulation

To prepare blank protein ink formulation, 3 mL of 12% w/v GelMA was prepared by dissolving lyophilized GelMA in DI water and mixed using a magnetic stirrer at 60℃. Once completely mixed, the GelMA solution was equally distributed into three separate vials, and the temperature was lowered to 40℃. To each vial, 1 mL of either 1%, 2%, or 4% w/v Laponite XLG solution was added and mixed with the stirrer. The resulting final compositions were 6% GelMA with 0.5%, 1%, or 2% Laponite XLG. Following the same printing procedure described above for polysaccharide ink formulation, blank grid structures were printed and compared, leading to a down-selection of 6% GelMA with 1% Laponite XLG formulation. Similarly, for biosensing ink formulation, 44 mg of reference microparticles (3X pre-washed with DI water) were first mixed with 2% w/v Laponite XLG before addition to 1 mL volume of 12% GelMA at 40℃. The mixture was quickly filled in the syringe barrel and 3D printed to obtain the grid structures. Finally, printed structures were stored in a 10 mM TRIS buffer (7.2) supplemented with 10 mM CalCl2 at 4℃ until further tests were performed.

#### Synthetic ink formulation

Blank synthetic ink formulations were made by mixing 1mL of 40% w/v PEGDA solution premixed with 1.12% w/v Irgacure 2959, with 1mL of 1%, 2%, 4%, or 8% Laponite XLG solutions. This resulted in three formulations with different Laponite XLG concentrations but with a fixed concentration of 20% w/v PEGDA and 0.56% w/v photoinitiator. The formulations were 3D printed to form grid structures. Since PEGDA does not possess any shear-thinning ability on its own, it was only Laponite XLG concentrations that dictated the printability behavior. Structures were printed over glass slides and exposed to a UV lamp for 5 minutes over each side at 20 mW/cm^2^ dosage. We downselected the 20% PEGDA with 4% Laponite XLG formulation since it offered the best printability results at the lowest polymeric concentrations. A similar procedure was used to make biosensing synthetic ink formulation. First, 44 mg of reference microparticles were washed three times with DI water, the supernatant discarded and mixed with 1 mL of 40% w/v PEGDA (premixed with 1.12% Irgacure 2959) for 10 minutes, protected from light. The mixture was then gradually added with 1 mL of 8% Laponite XLG, mixed with a spatula, and vortexed for 30 seconds. The resulting ink formulation was then filled into a syringe barrel for 3D bioprinting grid structures on glass slides, followed by UV lamp exposure on each side for 5 minutes. Finally, printed structures were stored in a 10 mM TRIS buffer (7.2) supplemented with 10 mM CalCl2 at 4℃ until further tests were performed.

### In-vitro sensor characterization - oxygen sensitivity responses

Oxygen sensitivity of Reference MPs in various polymeric formulations from hydrogel slabs and 3D bioprinted grid structures were measured under *in vitro* conditions using a setup described in **Supplemental Figure S4A**. Briefly, circular discs [from slabs; 3 mm (dia) × 0.75 mm (h)] or cut strips [from grids; 3 mm x 800 μm(dia.)] were secured over an acrylic sheet holder with rubber cantilever retainers and housed inside a rectangular flow cell. The flow cell had optical ports on one side that allowed connection with external readers for optical interrogation and readout measurements. Sensors were exposed to a circulating 500 mL volume of TRIS buffer solution premixed with varying dissolved oxygen concentrations (0 to 257.9 µM), achieved by controlling the air-nitrogen mixture at predetermined ratios. Gases were sparged and stirred in a buffer inside a 37°C incubator. Under these conditions, sensor lifetimes were measured using a time-domain phosphorescence lifetime reader system (630 nm excitation and 800 nm emission wavelengths), and quenching data fit a Stern–Volmer model.

### In-vitro sensor characterization - glucose or lactate challenge responses

The characterization of glucose- and lactate-sensing responses involved securing sensor samples, either circular punches or cut strips, within the flow cell assembly, matching the dimensions used for oxygen sensitivity experiments. These sensors were subsequently exposed to a series of TRIS buffer solutions containing increasing concentrations of either dissolved glucose (0, 100, 200, 300, and 400 mg/dL) or dissolved lactate (0, 10, and 20 mg/dL). Two reservoir solutions were employed: a stock solution (400 mg/dL glucose or 20 mg/dL lactate in TRIS buffer) and a blank TRIS buffer solution. The desired glucose or lactate concentrations were achieved within the flow cell for specific durations by precisely controlling the flow rates of these solutions using connected pneumatic pumps. Phosphorescence lifetime measurements were then recorded from the glucose sensors, followed by the generation of calibration plots based on the collected data.

### Rheological Tests

A suite of rheological tests was performed on the ink samples using a TA Instruments Discovery HR-2 Rheometer equipped with a 20 mm parallel plate geometry.

#### Shear rate sweep tests

At room temperature, ink samples were subjected to increasing shear rates (8 s⁻¹ to 800 s⁻¹) to measure viscosity changes.

#### Oscillatory stress sweep tests

Performed at room temperature, ink samples were exposed to varying stress (0.1 Pa to 1000 Pa) with recorded storage and loss modulus values. Yield stress was calculated in TRIOS software as the onset value of the storage modulus curves.

#### Peak hold tests

Employed to assess ink material recoverability after MP addition. This three-step process involved: 1) holding the ink under low stress (2-40 Pa) for 60 seconds, 2) briefly applying constant stress (400-600 Pa) for 5 seconds, and 3) removing constant stress for 120 seconds. For GelMA-based ink samples, all tests were conducted at 37°C to account for their gelation temperature. A separate Peak Hold Test dedicated for GelMA based bio-inks were performed to mimic viscosity changes during bioprinting process involving varying shear rates and temperatures. Given three-steps were followed: 1) holding the ink under low stress (2-40 Pa) for 60 seconds at 37°C, 2) briefly applying constant stress (400-600 Pa) for 5 seconds at 37°C, and 3) removing constant stress for 180 seconds at 25°C.

### Histology procedure

To further establish the long term biocompatibility of the microspheres and their performance over time within the hydrogels, acellular constructs were printed and inserted subcutaneously just under the skin of a healthy, immunocompetent pig. This position allowed us to evaluate the biosensors performance over an extended period of three months by reading the sensors noninvasively through the skin, and let us evaluate the biocompatibility of the hydrogel at the conclusion of the experiment. Cells were excluded from these experiments as the presence of foreign cells in an in vivo setting could be expected to independently elicit an immune response.

A healthy 7.5 month old female Yorkshire cross pig was used for this experiment (Pig 14). A Yorkshire cross was chosen for this experiment because they are well known as a production breed with robust health and a gregarious nature. A female was used due to ease of handling after puberty, and because it allowed the pig to live with her sister (Pig 13) as a companion animal for the duration of the experiment. Both pigs lived together in semi-enclosed housing, with access to fans and heat lamps as weather required, as well as hay beds, a wading pool, and a spectrum of toys and snacks for enrichment. Animals were interacted with daily by research and veterinary staff to encourage sociability and ease of handling. The animal study was IACUC approved under AUP# IACUC 2021-0066 Reference Number: 137661.

The bioprinted biosensors were subcutaneously inserted into Pig 14 at 7.5 months of age. Prior to insertion, she was anesthetized using isoflurane and intubated for safety. She was set on a heating pad and kept partially under a towel to keep her warm. Insertion sites were chosen in varied locations on the animal, from the thick shoulder skin to the relatively thin flank. Insertion sites were prepared first by closely trimming the hair, then cleaned with a mild detergent followed by isopropyl alcohol. A small marking was tattooed around each insertion site because the small inserted scaffolds were otherwise invisible under the skin. Biosensors were loaded into 16-gauge needles, which were subcutaneously inserted to a target depth of around 2 mm. A steel dowel rod was inserted through the needle to hold the insert in place as the needle and dowel rod were both steadily withdrawn. After the procedure, she was kept under observation until anesthesia had fully worn off, then returned to her pen and sister.

To conduct readings evaluating the construct’s performance, Pig 14 was led into an wheeled transport cage in her pen. She was otherwise unrestrained, as the procedure was painless and noninvasive. Pig 14 typically napped during readings but was allowed out regularly to stretch her legs.

Pig 14 was humanely euthanized at the conclusion of the experiment, 3 months after the insertion procedure. Her hide was skinned off and stored in neutral buffered formalin for at least a day. The insertion sites were then sectioned off using the tattoos as a guide, then the reader was used to precisely locate the sensor within the tissue. a square inch tissue surrounding the strongest sensor reading was cut out, then carefully sectioned and paraffin embedded, then stained with hematoxylin and eosin. The slides were prepared and examined by independent veterinary pathologists to evaluate device presence, device state, skin zone, host interface, and healing response.

### Cell culture procedures

Induced pluripotent cell derived mesenchymal stem cells (ihMSCs) were provided by Dr. Fei Liu (Texas A&M College of Medicine) and cultured in complete culture medium composed of alpha mem essential media, 100 U/ml penicillin/streptomycin, 4 mM L-Glutamine and 10% v/v fetal bovine serum. Media exchanges were performed every 2 days. When the cells reached approximately 70% confluency they were passaged by treatment with 0.25% trypsin, 0.1% EDTA (Corning) at 37°C for 5 min. The trypsin was deactivated with CCM, collected and centrifuged at 500 g for 5 min. Cells were collected and counted using a hemacytometer prior to seeding sensing experiments.

Ovarian cancer cells (OVCAR-3) were generously gifted by Dr. Anathakrishnan S. Jeevarathinam (MD Anderson Cancer Center). Cells were cultured in complete culture media composed of DMEM high glucose, 10% dialyzed FBS, 1X penicillin/streptomycin antibiotics. Similarly, media exchanges were performed every 2 days, and upon reaching 70% of cell confluency, cells were passaged using 0.25% trypsin, 0.1% EDTA at 37C for 5 min. Following trypsin deactivation, cells were centrifuged at 400 g for 4 mins, resuspended and counted prior to seeding sensing experiments.

### Real-Time Cell Culture Monitoring with biosensors

To evaluate the real-time functionality of biosensing bioink formulations, we separately incubated multianalyte biosensors with ihMSCs (non-transformed) and OVCAR-3 (transformed) cell lines. GelMA-based ink formulation was utilized to fabricate glucose-, lactate-, oxygen-, and temperature-sensing hydrogel slabs, circular discs [4 mm (dia) x 0.75 mm (h)] were punched out from each slabs, followed by sensor sterilization through Ebeam irradiation at National Center for Electron Beam Research, Texas A&M University (<25 KGy dose). Later, sensor discs were first strategically positioned over a 60 mm Petri dish plate, fixed by a rubber cantilever, and then Petri dish plate was fixed over a 3D- printed PETG scaffold. The scaffold firmly fixed the Petri Dish over its top surface. In contrast, the PETG scaffold’s underneath surface was used to attach optic fiber cables - directly interrogating the sensing spots from the bottom of the Petri dish plate. Finally, the other end of the optic fiber was interfaced with the 1D readers, allowing seamless and continuous interrogation of multi-sensors in the Petri dish by 1D readers. The entire assembly was placed over a stirrer, inside an incubator at 37℃ with 5% CO2.

A general procedure was followed to culture and monitor both cell lines where all four sensors were incubated in a single Petri dish plate along with their respective 5mL of complete media (with glucose) for 2-6 hours under stirring (speed 6). Later, cells were added to the plate at 0.2-0.4 million cells/ml of seeding density, and stirring was halted for 6-18 hrs. to allow cell attachment to the plate surface. The next day, microscopy images were acquired, media was replenished with a fresh complete media of the same volume, and stirring was initiated (speed 6). The cells were left to grow for the next 3-5 days, without changing any media, and glucose and lactate sensors were monitored for glucose level reduction and lactate production/surges. Once sensor response changes were observed, media was replaced with a DMEM incomplete media (without glucose) for a few hours, followed by the next exchange with respective complete media (with glucose) to validate glucose/lactate sensor activities. Before each media replenishment, a small volume (200µL) of media aliquots was collected for YSI measurements.

### Cytocompatibility Assessments

To determine the cell viability of ihMSCs and OVCAR-3 cells growing in well plates with multi-sensors, a live/dead staining was performed. The cells were washed 2 times cells with PBS. Then they were treated with PBS containing 5 µM calcein AM and 4 µM ethidium bromide for 30 min. Cell microscopy was performed (BiotTek, Cytation 5) where green fluorescence indicated live cell esterase activity and red fluorescence indicated ethidium bromide binding to the DNA in nonviable cells.

### Statistical Analyses

Statistical analyses were performed in GraphPad Prism 8 for comparison within and between groups. T-tests and, where appropriate, one-way ANOVA with appropriate post-hoc tests were performed. Statistical significance was set at p<0.05.

## Supporting Information

Supporting Information is available from the Wiley Online Library or from the author.

## Acknowledgments

The authors thank Dr. Staci Horn, Dr. Fred Club with histological analyses, and Lainey Streetman for providing help with rheology data collection. Some of the images in the article were created with Biorender. Research reported in this paper was supported by National Institutes of Health under award number R01EB024601.

